# Validation of a novel multiplex real-time PCR assay for *Trypanosoma cruzi* detection and quantification in açai pulp

**DOI:** 10.1101/2020.10.15.340984

**Authors:** Paula Finamore-Araujo, Amanda Faier-Pereira, Carlos Ramon do Nascimento Brito, Eldrinei Gomes Peres, Klenicy Kazumy de Lima Yamaguchi, Renata Trotta Barroso Ferreira, Otacilio Cruz Moreira

**Author notes:** Corresponding author. (OCM).

## Abstract

In Brazil, orally acquired *T. cruzi* infection has become the most relevant transmission mechanisms from public health perspective. Around 70% of new Chagas disease cases have been associated with consumption of contaminated food or beverages. Açai (*Euterpe oleracea* and *Euterpe precatoria*) is currently one of the most commercialized Amazonian fruits in the Brazilian and international markets. Therefore, it has become important to incorporate in the production process some procedures to measure out effective hygiene and product quality control required by global market. Molecular methods have been developed for rapid detection and quantification of *T. cruzi* DNA in several biological samples, including food matrices, for epidemiological investigation of Chagas disease and food quality control. However, a high-performance molecular methodology since DNA extraction until detection and quantification of *T. cruzi* DNA in açai berry pulp is still needed. Herein, a simple DNA extraction methodology was standardized from the supernatant of açai berry pulp stabilized in a Lysis buffer. In addition, a multiplex real time qPCR assay, targeting *T. cruzi* DNA and an Exogenous Internal Positive Control was developed and validated, using reference from all *T. cruzi* DTUs and commercial samples of açai pulp, from an endemic municipality with previous history of oral Chagas disease outbreak. Thus, a high-sensitivity qPCR assay, that could detect up to 0.01 parasite equivalents/mL in açai, was reached. As of the 45 commercial samples analyzed, 9 (20%) were positive for *T. cruzi*. This high-sensitive, fast and easy-to-use molecular assay is compatible with most of the laboratories involved in the investigations of oral Chagas disease outbreaks, representing an important tool to the epidemiology, control and surveillance of Chagas disease.

**Author Summary:** Oral transmission of Chagas disease has acquired an increasingly importance on the disease epidemiology. Most of the orally acquired Chagas Disease cases are related to the consumption of fresh foods or drinks, as sugar cane juice, açai berry pulp and bacaba wine, contaminated with triatomines or its feces. In Brazil, it has recently caused numerous outbreaks and has been linked to unusually severe acute infections. So far, the evaluation of the potential for oral transmission of Chagas disease through the consumption of açai-based products is mostly determined by clinical or parasitological methods. Despite the recent advances, a highly sensitive, reproductible and properly validated real time PCR assay for the molecular diagnostic of *T. cruzi* in açai pulp samples is still missing. Herein, a simple and reproducible multiplex real-time PCR assay was developed to the detection and quantification of *T. cruzi* DNA in açai pulp samples. This methodology, that includes a simple step for sample stabilization and DNA extraction based on silica-membrane spin columns, can be useful for analyzing orally transmitted acute Chagas disease outbreaks.

## Introduction

Chagas disease is a neglected tropical illness, caused by the flagellated and heteroxene protozoan *Trypanosoma cruzi*. Although it is considered endemic in 21 countries in America, mainly affecting Latin America, Chagas disease has now spread to previously non-endemic areas, due to the increased population migration between Latin America and the rest of the world [1 – 3]. As of today, it is estimated that six to seven million people are infected by *T. cruzi*, mostly in the endemic areas, and approximately 75 million people are at risk of infection [4]. *T. cruzi* is represented by a set of sub-populations, comprising isolates and strains, which alternates between mammalian hosts and insect vectors, with high genetic variability and notable heterogeneity of clinical behavior and parasite-host relationship [5 – 11]. Currently, seven genotypes, or Discrete Typing Units (DTUs) are recognized: TcI-TcVI, and Tcbat, this being last reported as TcVII [10, 11].

While Chagas disease is still often associated as a vectorborne disease, its transmission can occur in other different routes besides vectorial such as blood transfusions, organs transplantation, congenital, laboratory accidents and oral transmission [12 – 16]. However, foodborne outbreaks of Chagas disease seem to be importantly increasing through Latin America [17 – 19]. Oral transmission can occur due to the consumption of complete triatomines or its feces, containing metacyclic trypomastigotes, which is inadvertently processed with food, especially fruits preparations [20 – 26].

In Brazil, the oral route of *T. cruzi* infection has become one of the most relevant transmission mechanisms from the public health perspective. Between the years 2000 and 2011, 1252 cases of acute Chagas disease were reported and, in these, 70% have been associated with consumption of contaminated food or beverages [18, 26, 27]. Some of these orally acquired Chagas Disease are related to the consumption of fresh foods or drinks, as sugar cane juice, açai berry (*Euterpe oleracea*) and bacaba wine (*Oenocarpus bacaba*) [18, 19]. Among the brazilian outbreaks, it is worth mentioning a suspected *T. cruzi* oral transmission through açai juice in the Northern Brazil, where cases of this transmission route show signs of increase. The outbreak occurred in 2008 at Coari city, in the interior of the Amazonas state, where 25 people got infected by *T. cruzi* [25, 28]. Other countries around Latin America, suchlike Argentina, Bolivia, Colombia, Ecuador, and Venezuela, also have reported multiple Chagas disease outbreaks acquired orally [19]. The largest outbreak described so far occurred in Venezuela, in 2007, affecting 103 people, adults and children from a school in Caracas, that got infected by *T. cruzi* after the consumption of contaminated guava juice [24].

The açaizeiro is one of the most important socioeconomically palm tree that occurs especially throughout the Amazon region and is particularly abundant in the Eastern Amazon [29, 30]. Three species are popularly known as açai, *Euterpe edulis, E. oleraceae and E. precatoria*, however, only the last two have agro-industrial interest [31]. Both species have been shown antioxidant and anti-inflammatory properties with bioactive compounds as anthocyanins, flavonoids and phenolics, more than other fruits, such as grapes, blackberries, blueberries and strawberries [32. 33]. Its development takes place in floodplains and in swampland and flooded soils [34]. Fruiting occurs throughout the year, with the highest production of the açai berry occurring during the period of July to December [34, 35]. Most of the açai is derived from the extractive activity, which is the main source of income for the riverine population of the Amazon Basin. In the North of Brazil, mainly, most of the production is still consumed by the local population, in which açai is traditionally ingested "*in natura*", especially in Para and amazon states, the biggest producers and consumer of açai around the world [36]. Furthermore, in the last ten years, there has been an important economic increase, nationally and internationally, in açai-based drinks commercialization. Açai is currently one of the most commercialized Amazonian fruits, not only in the Brazilian market, but also at an international level, like in United States, that consumes most of all açai that is exported, Asia and Europe [30, 37]. Even so, the national and international markets still have potential for considerable expansion of açai commercialization. Therefore, it has become important to incorporate in the production process some procedures to measure out effective hygiene and product quality control required by global market [38 – 40].

Until now, the evaluation of the potential for oral transmission of Chagas disease through the consumption of açai-based products is determined by clinical or parasitological investigations and are based on traditional methods, such as culture and microscopic observations [15,19]. However, culture is a laborious method associated with risk of contamination [41] and microscopic analysis is a toilsome and time-consuming procedure, besides being difficult to detect the parasite during the examination, due to açai’s characteristic dark color [42]. Thus, diagnosis using molecular or serological tests in patients can assist outbreaks investigations and overcome these limitations [19]. Molecular methods based on PCR have been developed for the rapid detection and quantification of *T. cruzi* DNA in several biological samples [43 – 49]. More recently it has also begun to be used to test food matrices, as a powerful tool in the epidemiological investigation of Chagas disease [26, 41, 50 – 52]. Moreover, the use of PCR for the detection of *T. cruzi* DNA has already been described in literature, since several studies demonstrate that this methodology presents greater sensitivity and specificity in the face of conventional parasitological methods [53]. Nevertheless, it is still demanded the development of methodologies capable to detect *T. cruzi* in food samples and, thereby, plan strategies to get a better understanding of oral transmission and to assure the quality of the products thar are being commercialized [34, 51].

Based on that, the aim of this work was the development and validation of a rapid molecular methodology, simple and reproducible, based on real-time PCR to the detection and quantification of *T. cruzi* DNA in açai samples. This proposed methodology, that includes a simple step for sample stabilization and DNA extraction based on silica-membrane spin columns, can be useful for analyzing orally transmitted acute Chagas disease outbreaks.

## Results

The present work aimed to develop an appropriate molecular methodology based on a real-time PCR assay to rapidly assess, with reproducible protocols, the presence of *T. cruzi* DNA in açai pulp samples, facilitating the food quality control and investigations of Chagas disease oral outbreaks involving açai-based products. Therefore, an efficient DNA extraction method for the isolation of *T. cruzi* DNA from complex food matrix as açai were also developed and tested. To increase the sensitivity of the methodology, artificially contaminated açai samples were submitted to a pre-lysis stage, through the addition of a lysis buffer (Guanidine-HCl 6M/EDTA 0.2M pH 8.0), followed by a centrifugation step as described in material and methods section. To monitor the entire procedure concerning the stability of DNA and its loss during sample processing, a normalized amount of an exogenous internal positive control DNA (EXO-IPC DNA) was added to each GEA immediately before DNA extraction with the High Pure PCR Template Preparation kit (Roche Life Science, Mannheim, Germany). Then, through qPCR assays, EXO-IPC DNA amplification from GEA samples with low *T. cruzi* concentrations was assessed (Fig 1).

**Fig 1.**
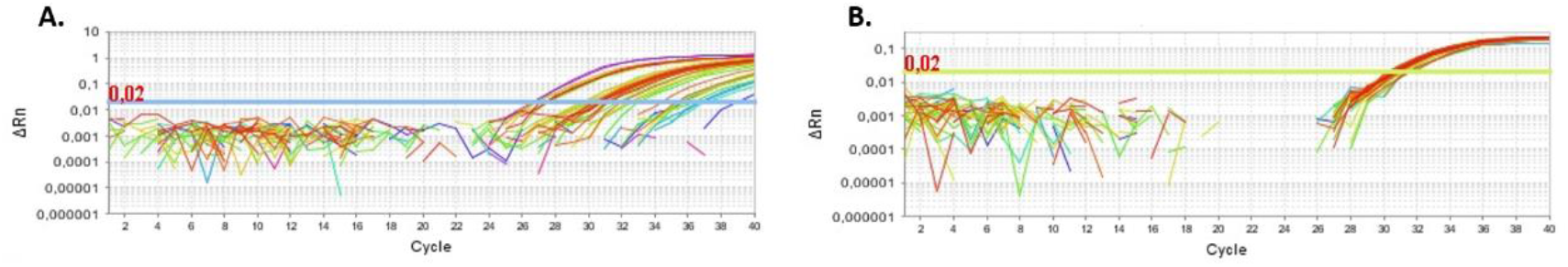
Representative amplification plots, targeting T. cruzi nuclear satellite DNA (A) and EXO-IPC DNA (B). DNA samples were extracted from GEA spiked with EXO-IPC synthetic DNA, using a silica-column based commercial kit.

In order to evaluate the analytical sensitivity of the multiplex real time PCR assay, GEA samples were spiked with *T. cruzi* and serially-diluted to reach from 10 to 0.01 parasite equivalents/mL, prior the supernatant isolation and DNA extraction. In addition, 300 μL supernatant aliquots were spiked with EXO-IPC DNA, and satDNA and Exo-IPC amplifications were monitored trough the Ct values at the real time PCR assay. As showed in Table 1, it is possible to observe the satDNA amplification in all concentrations tested, with Ct values from 27.32±0.34 to 36.82±1.3, reaching a sensitivity of 0.01 *T. cruzi* equivalents/mL. Furthermore, the Ct values for the Exo-IPC varies only from 30.50±0.12 to 30.92±0.44, regardless the *T. cruzi* concentration at the samples, in the range tested.

**Table 1.**
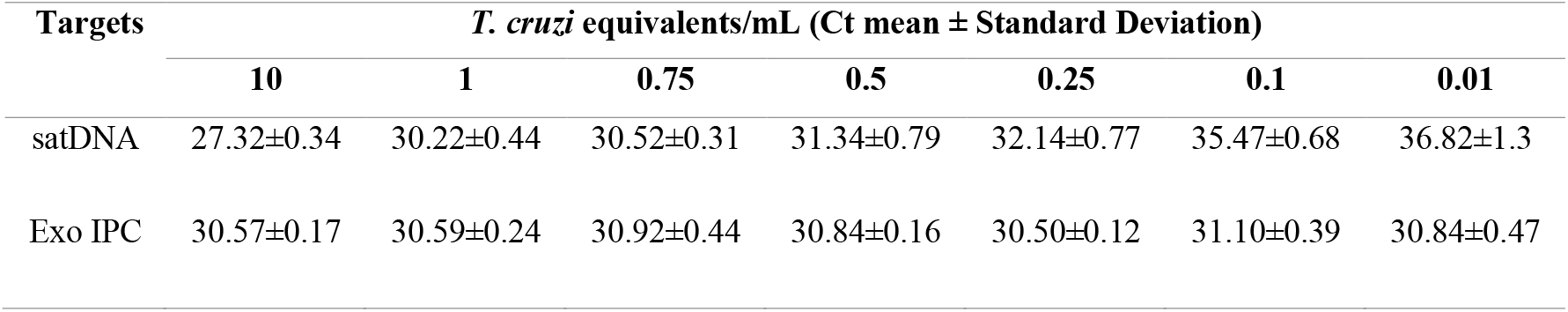
Analytical sensitivity for the satDNA detection in GEA samples spiked with *T. cruzi* and Exo-IPC DNA. Assays were tested with genomic DNA extracted from GEA samples with different *T. cruzi* (Dm28c) concentrations, ranging from 10 to 0.01 *T. cruzi* equivalents/mL. All GEA supernatant samples were spiked with 2 μL EXO-IPC DNA prior DNA extraction. Results are shown as Ct mean ± SD for satDNA and Exo-IPC at the multiplex real time PCR assay.

Following the analytical validation of the multiplex real time PCR assay, the reportable range for the *T. cruzi* load quantification in açai pulp samples was determined. It was possible to observe that the *T. cruzi* DNA detection presented an improved linearity (r^2^=0.99) in the range from 10^6^ to 1 parasite equivalents/mL, to the serial dilution of DNA extracted from GEA spiked with a pool of *T. cruzi* epimastigotes (Fig 2). Under these conditions, it was possible to obtain a PCR efficiency of 89.65% to the amplification of the satDNA target. In addition, no Ct outlier was observed for the EXO-IPC target in any point of the standard curve (data not shown).

**Fig 2.**
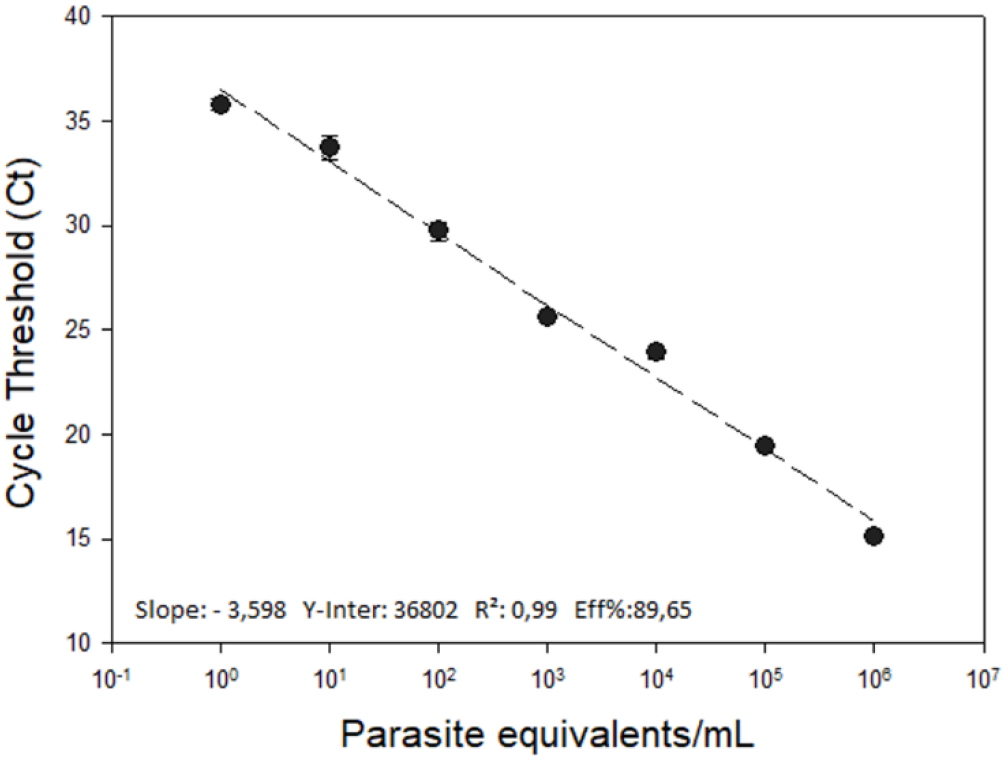
Reportable range for the quantification of *T. cruzi* in açai samples by real-time qPCR. A ten-fold serial dilution of DNA extracted from GEA spiked with *T. cruzi* was used to generate the standard curve for sat-DNA target, ranging from 10^6^ to 1 Par. Eq./mL. The bottom of the graphic shows the standard curve parameters of the assay.

Thereafter, inclusivity and exclusivity assays were performed. For the inclusivity assay, the amplification of *T. cruzi* satDNA from strains or clones belonging to TcI to TcVI was evaluated in GEA samples spiked with the parasites. Table 2 shows that the multiplex qPCR assay could detect all the six *T. cruzi* DTUs, from 10^4^ to 0.1 Par. Eq./mL, with Ct values varying from 20.37±1.25 (to TcV, at 10^4^ Par. Eq/mL) to 42.80 (to TcVI, at 0.1 Par. Eq/mL).

**Table 2.**
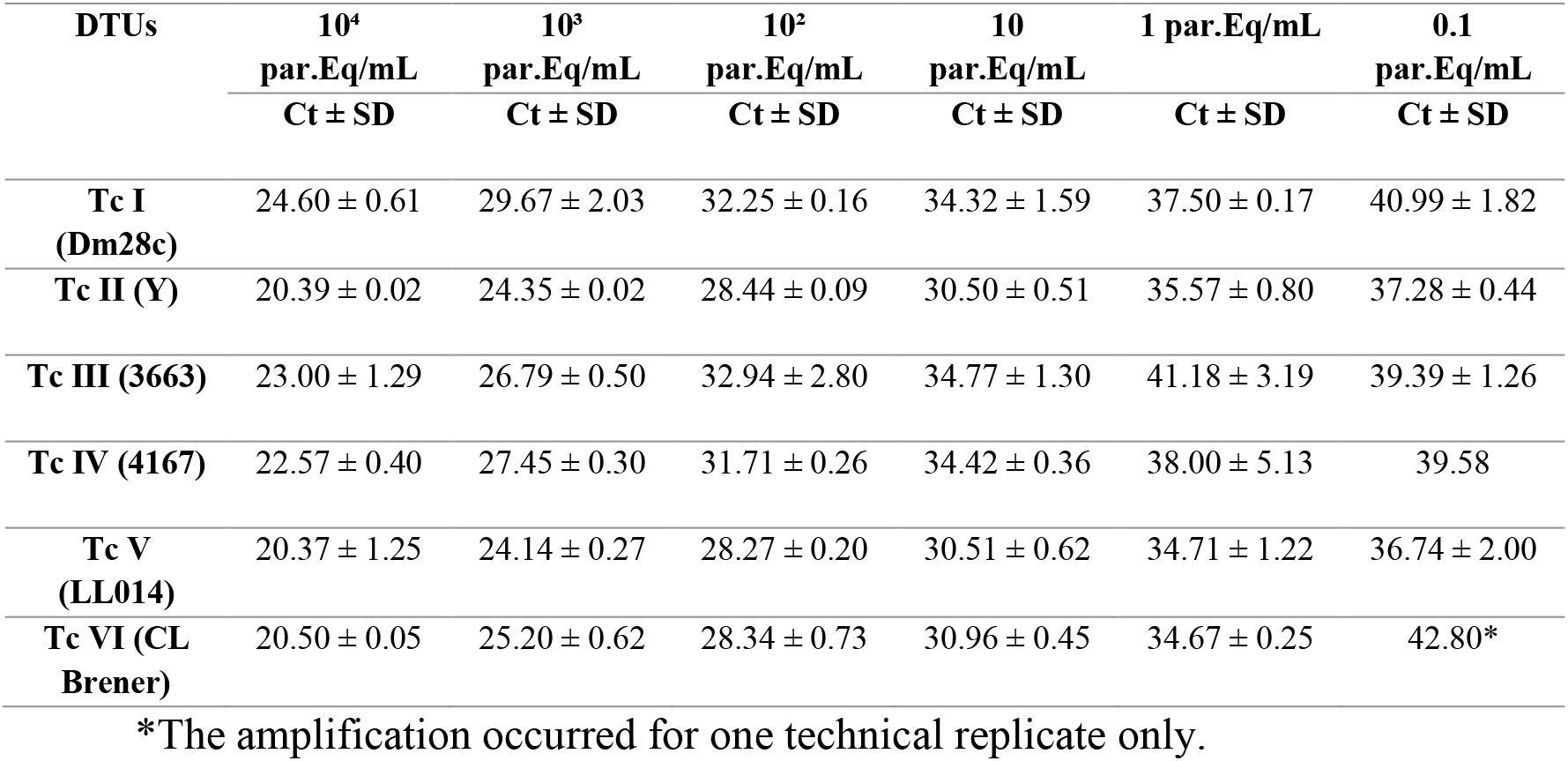
Inclusivity assay for *T. cruzi* DTUs. Assays were tested with genomic DNA obtained from a panel of GEA samples spiked with *T. cruzi* from six different DTUs. DNA concentrations ranged from 10^4^ to 10^−1^ parasites equivalents/mL. Results are shown as Ct mean ± SD obtained from duplicates of each DNA concentration.

Regarding the exclusivity assay, GEA samples were spiked with different species from the Trypanosomatidae family (*Leishmania (L.) amazonensis*, *Leishmania (V.) braziliensis*, *Crithidia* sp., *Herpetomonas* sp. and *Trypanosoma rangeli*) and the cross-amplification in the multiplex real time PCR assay targeting satDNA and EXO-IPC was investigated. It is possible to observe in Table 3 that DNA of other trypanosomatids, as *Leishmania (L.) amazonensis*, *L (V.) braziliensis*, *Crithidia* sp. and *Herpetomonas* sp., showed no amplification in any concentration tested. However, when we tested different concentrations of DNA extracted from GEA spiked with *T. rangeli*, qPCR was positive for all concentration tested, from 10^4^ parasites equivalents/mL to 0.1 par. Eq./mL.

**Table 3.** Exclusivity assay with other Trypanosomatids. Assays were tested with serial dilutions of purified DNAs from different species of Trypanosomatids that ranged from 10^4^ to 0.1 parasites equivalents/mL. Results are shown as Ct mean ± SD obtained from duplicates of each DNA concentration.

| Samples | 10<sup>4</sup><br>par.Eq/mL<br>Ct ± SD | 10<sup>3</sup><br>par.Eq/mL<br>Ct ± SD | 10<sup>2</sup><br>par.Eq/mL<br>Ct ± SD | 10<br>par.Eq/mL<br>Ct ± SD | 1<br>par.Eq/mL<br>Ct ± SD | 0.1<br>par.Eq/mL<br>Ct ± SD |
| --- | --- | --- | --- | --- | --- | --- |
| *L. (L.) amazonensis* | NA | NA | NA | NA | NA | NA |
| *L. (V.) brasiliensis* | NA | NA | NA | NA | NA | NA |
| *Crithidia* sp. | NA | NA | NA | NA | NA | NA |
| *Herpetomonas* sp. | NA | NA | NA | NA | NA | NA |
| *T. rangeli* | 22.95 ± 0.58 | 27.83 ± 0.62 | 31.18 ± 0.30 | 32.95 ± 0.20 | 33.94 ± 0.64 | 38.91 ± 0.60 |

NA: No amplification

The last step of this study was to perform the clinical validation of the multiplex qPCR assay. Therefore, 45 samples of açai pulp were collected from different street markets in the city of Coari (Amazonas State, Brazil) and investigated for the presence of *T. cruzi* by direct observation at a light microscope and by the qPCR assay. DNA obtained from GEA samples, following the adapted protocol for DNA purification directly from the supernatant, was used to assess the performance of the qPCR multiplex methodology standardized in this work. No *T. cruzi* could be detected by microscopy in the açai pulp samples Nevertheless, it was possible to detect and quantify *T. cruzi* in 9 samples (20%) by the multiplex qPCR, with parasite loads ranging from 0.002 to 19,05 par. Eq./mL, as showed in Table 4. In addition, all samples amplified to the Exo-IPC target, with Cts from 28.51 ± 0.75 to 34.94 ± 0.48, validating the true-negative samples.

**Table 4.**
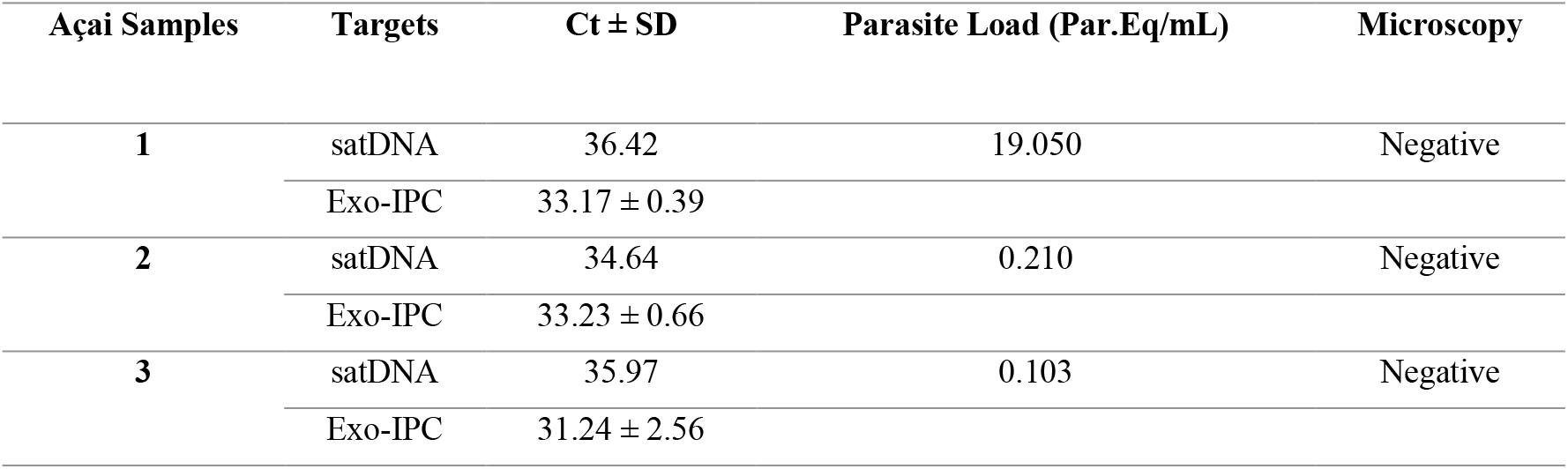
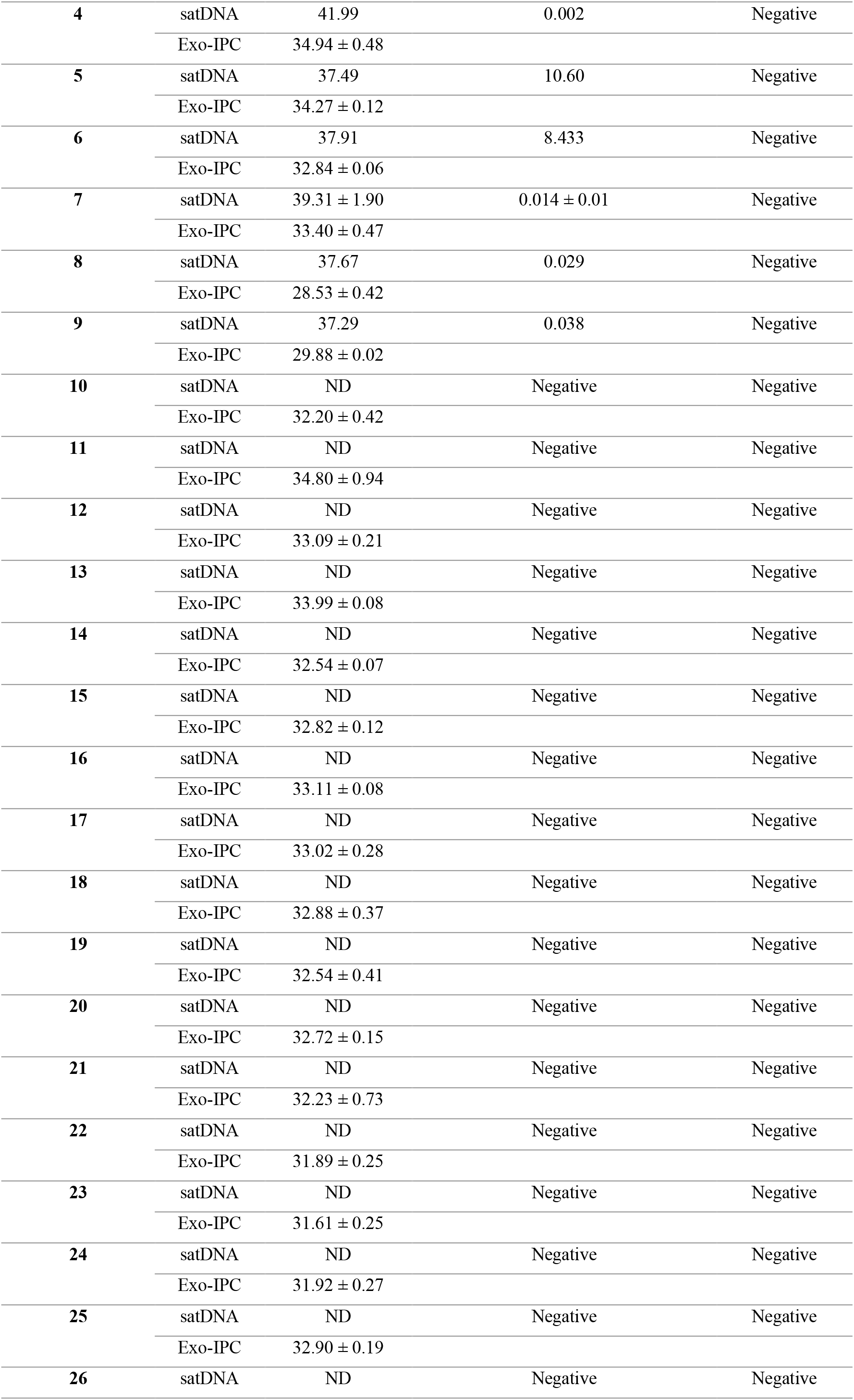
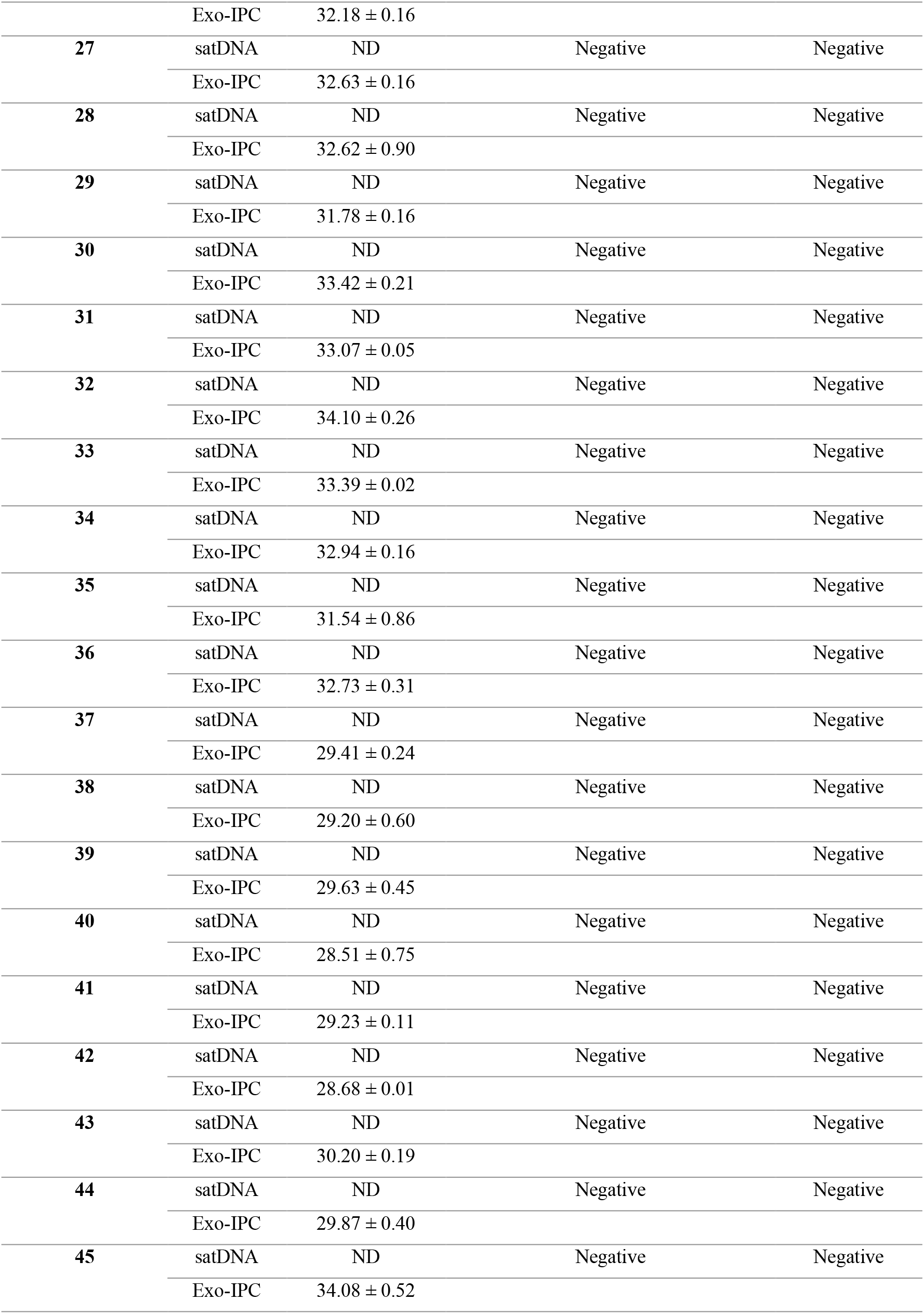
Validation of the multiplex real time qPCR assay. Fourty-five açai pulp samples were obtained from different points of sale at the Coari city (Amazonas State, Brazil). *T. cruzi* was detected and quantified using primers targeting satDNA and exo-IPC, in the multiplex qPCR assay. Results are indicated as Ct mean ± SD and parasite load is shown as parasite equivalents/mL.

| Açai Samples | Targets | Ct ± SD | Parasite Load (Par.Eq/mL) | Microscopy |
| --- | --- | --- | --- | --- |
| 1 | satDNA | 36.42 | 19.050 | Negative |
|  | Exo-IPC | 33.17 ± 0.39 |  |  |
| 2 | satDNA | 34.64 | 0.210 | Negative |
|  | Exo-IPC | 33.23 ± 0.66 |  |  |
| 3 | satDNA | 35.97 | 0.103 | Negative |
|  | Exo-IPC | 31.24 ± 2.56 |  |  |
| **4** | satDNA | 41.99 | 0.002 | Negative |
|  | Exo-IPC | 34.94 ± 0.48 |  |  |
| **5** | satDNA | 37.49 | 10.60 | Negative |
|  | Exo-IPC | 34.27 ± 0.12 |  |  |
| **6** | satDNA | 37.91 | 8.433 | Negative |
|  | Exo-IPC | 32.84 ± 0.06 |  |  |
| **7** | satDNA | 39.31 ± 1.90 | 0.014 ± 0.01 | Negative |
|  | Exo-IPC | 33.40 ± 0.47 |  |  |
| **8** | satDNA | 37.67 | 0.029 | Negative |
|  | Exo-IPC | 28.53 ± 0.42 |  |  |
| **9** | satDNA | 37.29 | 0.038 | Negative |
|  | Exo-IPC | 29.88 ± 0.02 |  |  |
| **10** | satDNA | ND | Negative | Negative |
|  | Exo-IPC | 32.20 ± 0.42 |  |  |
| **11** | satDNA | ND | Negative | Negative |
|  | Exo-IPC | 34.80 ± 0.94 |  |  |
| **12** | satDNA | ND | Negative | Negative |
|  | Exo-IPC | 33.09 ± 0.21 |  |  |
| **13** | satDNA | ND | Negative | Negative |
|  | Exo-IPC | 33.99 ± 0.08 |  |  |
| **14** | satDNA | ND | Negative | Negative |
|  | Exo-IPC | 32.54 ± 0.07 |  |  |
| **15** | satDNA | ND | Negative | Negative |
|  | Exo-IPC | 32.82 ± 0.12 |  |  |
| **16** | satDNA | ND | Negative | Negative |
|  | Exo-IPC | 33.11 ± 0.08 |  |  |
| **17** | satDNA | ND | Negative | Negative |
|  | Exo-IPC | 33.02 ± 0.28 |  |  |
| **18** | satDNA | ND | Negative | Negative |
|  | Exo-IPC | 32.88 ± 0.37 |  |  |
| **19** | satDNA | ND | Negative | Negative |
|  | Exo-IPC | 32.54 ± 0.41 |  |  |
| **20** | satDNA | ND | Negative | Negative |
|  | Exo-IPC | 32.72 ± 0.15 |  |  |
| **21** | satDNA | ND | Negative | Negative |
|  | Exo-IPC | 32.23 ± 0.73 |  |  |
| **22** | satDNA | ND | Negative | Negative |
|  | Exo-IPC | 31.89 ± 0.25 |  |  |
| **23** | satDNA | ND | Negative | Negative |
|  | Exo-IPC | 31.61 ± 0.25 |  |  |
| **24** | satDNA | ND | Negative | Negative |
|  | Exo-IPC | 31.92 ± 0.27 |  |  |
| **25** | satDNA | ND | Negative | Negative |
|  | Exo-IPC | 32.90 ± 0.19 |  |  |
| **26** | satDNA | ND | Negative | Negative |
|  | Exo-IPC | 32.18 ± 0.16 |  |  |
| **27** | satDNA | ND | Negative | Negative |
|  | Exo-IPC | 32.63 ± 0.16 |  |  |
| **28** | satDNA | ND | Negative | Negative |
|  | Exo-IPC | 32.62 ± 0.90 |  |  |
| **29** | satDNA | ND | Negative | Negative |
|  | Exo-IPC | 31.78 ± 0.16 |  |  |
| **30** | satDNA | ND | Negative | Negative |
|  | Exo-IPC | 33.42 ± 0.21 |  |  |
| **31** | satDNA | ND | Negative | Negative |
|  | Exo-IPC | 33.07 ± 0.05 |  |  |
| **32** | satDNA | ND | Negative | Negative |
|  | Exo-IPC | 34.10 ± 0.26 |  |  |
| **33** | satDNA | ND | Negative | Negative |
|  | Exo-IPC | 33.39 ± 0.02 |  |  |
| **34** | satDNA | ND | Negative | Negative |
|  | Exo-IPC | 32.94 ± 0.16 |  |  |
| **35** | satDNA | ND | Negative | Negative |
|  | Exo-IPC | 31.54 ± 0.86 |  |  |
| **36** | satDNA | ND | Negative | Negative |
|  | Exo-IPC | 32.73 ± 0.31 |  |  |
| **37** | satDNA | ND | Negative | Negative |
|  | Exo-IPC | 29.41 ± 0.24 |  |  |
| **38** | satDNA | ND | Negative | Negative |
|  | Exo-IPC | 29.20 ± 0.60 |  |  |
| **39** | satDNA | ND | Negative | Negative |
|  | Exo-IPC | 29.63 ± 0.45 |  |  |
| **40** | satDNA | ND | Negative | Negative |
|  | Exo-IPC | 28.51 ± 0.75 |  |  |
| **41** | satDNA | ND | Negative | Negative |
|  | Exo-IPC | 29.23 ± 0.11 |  |  |
| **42** | satDNA | ND | Negative | Negative |
|  | Exo-IPC | 28.68 ± 0.01 |  |  |
| **43** | satDNA | ND | Negative | Negative |
|  | Exo-IPC | 30.20 ± 0.19 |  |  |
| **44** | satDNA | ND | Negative | Negative |
|  | Exo-IPC | 29.87 ± 0.40 |  |  |
| **45** | satDNA | ND | Negative | Negative |
|  | Exo-IPC | 34.08 ± 0.52 |  |  |

## Discussion

In the early 1990s, there was a milestone in the control of Chagas disease in South America. Countries in the Southern Cone (Argentina, Bolivia, Brazil, Chile, Paraguay, and Uruguay) adopted measures to control the vector and to screen and test blood banks [55, 56]. The implementation of programs for the triatomine elimination in Latin American countries has resulted in the control of the transmission in several endemic areas by its main vector, and a significant decrease in the incidence of new cases [1, 57 – 60]. Nevertheless, the possibility of vector transmission still prevails since *T. cruzi* circulates among other triatomines species and several small mammals in sylvatic environments in which there is human activity [11, 18, 59, 60]. In addition, several outbreaks of oral infection in endemic areas are generally related to the consumption of food contaminated with infected triatomines, or their feces, and with infected secretions of mammals [22, 23, 24, 25,61, 62].

*T. cruzi* transmission via the oral route is not a recent event, since it represents the main route of contamination between vectors and animals and also is one of the main mechanisms of parasite dispersion among mammals [11, 18]. Despite that, only after 2004 foodborne outbreaks of Chagas disease became an event most frequently discussed and investigated [63]. Originally sporadic reported, orally acquired Chagas disease seem to be increasing among populations in several Brazilian states [17–19] and in other endemic Latin American countries as well [19, 24, 62, 64]. However, in the Northern region of Brazil, the oral transmission poses significant importance, mainly due to the daily diet based on the consumption of food *in natura*, which means unpasteurized homemade or artisan fresh food [19, 22]. Regarding the food ingested, açai, which is macerated in order to produce a paste or drink, has been identified as the most frequent food involved in cases of orally acquired acute Chagas disease in Brazil [22, 63, 65]. Besides being important to the local culture, açai is also a fundamental source of income for the population of the Amazon Basin, contributes significantly to riverine and rural communities in Amazon, being a source of economic and social development in the region [36, 65], since it is one of the most commercialized Amazonian fruits, not only in the national market, but also at an international level [30, 37]. Even if industrialized and exported açai are supposed to be pasteurized, the majority of açai consumed by the population of Latin American countries is not treated or properly sanitized [41, 66] and the precise stage of food handling at which contamination occurs is not well enlightened [22].

*T. cruzi* oral transmission prevention has proved to be relatively difficult, giving rise to the public health system a new demand to face orally acquired Chagas disease. Although guidelines for minimizing contamination by microorganisms and parasites during the processing in the food chain have already been established [63, 67, 68], it is necessary to assess the quality of açai products that are sold and consumed by the population. Some studies that allow the molecular identification of the parasite in different food sources are already being developed and published [26, 41, 50–52, 63], however there is no official regulated method for the molecular detection of *T. cruzi* in açai pulp. Thus, it is necessary to develop an integrated, simple, and reproducible molecular methodology to supply this new demand of quality control and sanitary management required by local and global market for food safety.

The first step of the present work was to verify if the methodology chosen for DNA extraction was efficient to extract a high-quality genomic DNA from *T. cruzi* in açai samples. One of the biggest challenges for DNA extraction from parasites in such a complex food matrix lies on the presence of inhibitors that can be co-purified with the DNA during the extraction step and can reduce the efficiency of the PCR. Açai is composed of a significant portion of lipids, proteins, carbohydrates, soluble and non-soluble dietary fibers, fatty acids, a variety of minerals, and high antioxidant compound content, like anthocyanins, and phenolic compounds [33–35]. Presently, there are several methods available for DNA extraction from different sorts of samples and they can be based on commercial kits or can be in-house methods. Ferreira et al. [63] compared the quantity and quality of *T. cruzi* DNA isolated from artificially contaminated açai samples, whereby different DNA extraction methods were applied. In the study, the selected commercial kit proved not to be adequate for *T. cruzi* detection in food samples while one of the protocols that used CTAB (cationic hexadecyl trimethyl ammonium bromide) yielded satisfactory results, for use in PCR, regarding DNA purity and concentration. Souza-Godoi et al. [50] developed a methodology based on Real Time qPCR for *T. cruzi* detection and quantification in açai samples and a phenol-chloroform protocol was used for the DNA extraction. De Oliveira et al. [41] aimed to investigate, through DNA and RNA detection, prevention methods, as sanitization and thermal treatment, for orally transmitted Chagas disease. Therefore, a protocol using organic solvents was also used to perform DNA extraction. In contrast, when they compared the results through RT-PCR analysis, a commercial kit was used regarding mRNA isolation. Despite being low cost and offer a high concentration of DNA, in-house methods, such as phenol-chloroform and CTAB are often laborious and not suitable to assess large-scale analyzes, since they are not reproducible, simple, or rapid protocols. In relation to DNA extraction using a commercial kit, Mattos et al. [26], that developed a molecular methodology based on real-time qPCR, selected two different extraction kits. DNA was extracted using QIAamp DNA Mini Kit (Qiagen), and DNA from food samples was extracted using a specific kit for use in stool samples (QIAamp DNA Stool Mini Kit). Then, they observed that the stool kit permitted inhibitors removal, while others commercial kits did not show the same performance and, consequently, PCR was always negative even though acai samples were contaminated.

As a first attempt to increase the sensitivity of the methodology in this study, artificially contaminated açai samples were submitted to a pre-lysis stage, through mixing an equal volume of açai and 2X of a lysis buffer (Guanidine-HCl 6M/EDTA 0.2M pH 8.0) at room temperature. Guanidine-HCl can disrupt cells in addition to inhibiting nucleases, being a chaotropic salt commonly used for isolation of nucleic acids from cellular extracts. Thus, this reagent facilitates the preservation of nucleic acids in biological fluids [69, 70] and the sample transport from the field to the laboratory. Several previous studies related to molecular diagnosis of *T. cruzi* in blood samples have already incorporated this step into the methodology, due to its importance as a stabilizing buffer to the sample [46, 48, 70 – 75]. One question concerning DNA extraction based on silica-membrane spin columns is due to açai viscosity which can impair the procedure [26], mainly because of the silica-membrane clogging. Because of that, even though many commercial kits are available for extracting DNA from different matrices, only a limited number can be used for DNA purification from processed food products, which can make these kits even more expensive. Then, lysed GEA samples were submitted to a centrifugation step for parasite nucleic acid recovery and the supernatant was used for DNA extraction, using a commercial silica-column kit. Particularly, the kit used in this study has an inhibitor removal buffer that permits removing inhibitors residues that could have remained in açai samples.

Regarding Real Time qPCR, infection rates in complex food matrices may be underestimated since the chemical characteristics of açai may lead to a reduction of the PCR reaction due to the presence of contaminants and inhibitors [63]. The use of negative and positive controls is essential to ensure the reliability of the reaction, avoiding false-positive or false-negative results. However, especially in quantitative real-time qPCR assays, in addition to the negative and positive controls, it is also necessary to include an internal amplification control to monitor the reproducibility of DNA extraction and the absence of PCR inhibition (total or partial). In previous studies for the molecular diagnosis of Chagas disease, a target at the host DNA, such as RNAse P, or an exogenous DNA, were used as internal amplification controls, to avoid false negative results, which can occur when working with highly complex material [46, 48, 51, 73, 76, 77], and to correct and normalize DNA variations between samples. Ferreira et al. [51], in order to evaluate the amplifiability of DNA in a conventional PCR, employed a plant-specific primer pair [78], which encodes the ribulose 1,5-diphosphate carboxylase/oxygenase gene (*rbcL*) of the plant chloroplast. However, to date, published studies related to the development of a methodology based on Real-Time quantitative PCR, for the *T. cruzi* diagnosis in food samples, have not included an internal amplification control, despite using negative and positive controls in the reactions [26, 50, 52]. In our study, we used a synthetic DNA from a commercial kit (Applied Biosystems, Foster City, CA, USA) to monitor the efficiency of the DNA extraction and the absence of inhibitors at qPCR. The reproducibility of the qPCR was confirmed since there was no variation in Exo-IPC Ct values, regardless *T. cruzi* concentration at the samples. Besides that, our internal amplification control was amplified in all açai samples, validating the true-negative results.

In most outbreaks, the presence of *T. cruzi* in food is, normally, detected by traditional methods, such parasite isolation and microscopic investigation. And, although this practice presents results with high specificity, microscopic examination is a labor-intense, time consuming method and has low sensitivity, being minimally effective when only a few microorganisms exist in a sample [65, 79]. Besides that, Barbosa et al. [42] described about the non-visualization, by light microscopy, of the parasites directly from açai pulp, even with the addition of trypan blue to the samples. Namely, the characteristic dark color of the fruit, which is associated with high anthocyanin concentrations and a large amount of organic matter [34, 35], greatly limit the possibility of parasites detection during microscopic visualization. In this context, molecular methods can overcome these limitations regarding analysis directly from food, and provide specific diagnosis [26, 41, 50, 52, 63] and *T. cruzi* genotyping [51]. The qPCR developed in the present study showed improved sensitivity, in which was possible to detect the DNA corresponding to 0.01 *T. cruzi* equivalents/mL in the sample. Besides that, *T. cruzi* satDNA detection, with the set of primers and probes selected, presented an improved linearity and a PCR efficiency of 89.65%. Souza-Godoi et al. [50], who used phenol-chloroform protocol for DNA extraction, also obtained, with a SYBR-green qPCR methodology, a high linearity and an efficiency of 80.82% for the standard curve targeting *T. cruzi* satDNA. To assess the performance of the multiplex qPCR, targeting for *T. cruzi* sat-DNA, it was possible to observe that all the six *T. cruzi* DTUs could be detected in GEA samples, from 10^4^ to 0.1 Par. Eq./mL, in the inclusivity assay. Despite the remarkable inclusivity of the multiplex real time PCR assay, we could also observe that the Dm28c (TcI) presented higher Ct values for the satDNA target than the other DTUs, in general. This result corresponds with previous observations that the *T. cruzi* DTU I present a lower number of copies for the nuclear satellite DNA [73]. Regarding the exclusivity assay, it was possible to observe that DNA of other trypanosomatids species showed no cross-amplification in any concentration tested. However, when we tested different concentrations of *T. rangeli* DNA, qPCR was positive, confirming the previous observed cross-amplification of *T. rangeli* with this satDNA primers and probe set [48, 80]. This expected result did not impair the use of this methodology to detect and quantify *T. cruzi* in açai samples, in cases of Chagas disease oral outbreaks, since *T. rangeli* will not cause any symptoms in humans, in contrast to the acute infection triggered by *T. cruzi*. However, new molecular targets should be investigated in order to enhance the specificity for the *T. cruzi* parasite load quantification in açai samples.

In relation to the analysis of commercial açai-based samples, this work is, until now, the first one that has assessed, using a methodology based on Real-Time quantitative PCR, samples gathered from street markets of an endemic area for clinical validation. DNA of 45 samples of açai pulp from Coari city (Amazonas State, Brazil) were assayed according to our methodology. The qPCR screening of these GEA samples was shown to be more sensitive than microscopic examination, since the molecular method revealed a positivity of 20% (9/45), whereas all samples were negative for *T. cruzi* by direct observation at a light microscope. Before that, Ferreira et al. [51] determined, in food samples commercialized in Rio de Janeiro and Pará states, *T. cruzi* contamination rates and molecular characterization through conventional PCR and multilocus PCR analysis, respectively. For *T. cruzi* DNA detection, a PCR amplifying the telomeric region of gp85/sialidase superfamily was performed. And, although the set of primers [81] employed in the PCR showed high specificity, since there was no amplification for *T. rangeli* and other trypanosomatids genomic DNA, the chosen target had lower sensitivity. Interestingly, from the 140 samples of açai-based products they analyzed, 14 samples (10%) was positive for *T. cruzi* DNA and triatomine DNA was also detected in one of these 14 samples. A previous study [82] also found insect fragments in açai samples and these findings may reinforce the link between açai and the presence of infected vectors or mammals near of the outbreak locations.

Regarding the use of molecular methodologies as a diagnosis tool, *T. cruzi* DNA detection by itself does not mean the presence of viable parasite in the sample, once DNA molecule can remain stable even a little after parasite death [41, 83, 84]. Nevertheless, the detection of the parasite DNA can be important to identify problems related to good manufacturing practices throughout the production chain and in the epidemiological investigation of orally acquired Chagas disease. In this context, our results present a simple and rapid extraction protocol that provides a high-quality genomic DNA directly from açai sample, as well as a highly sensitive multiplex qPCR-based methodology, which includes a commercial exogenous internal positive control. The methodologies standardized herein can assist in the surveillance of commercialized açai-based products, in a large-scale basis, and can be a powerful tool for a better understanding about orally acquired Chagas disease and for strategies to assure the safety of açai, such as in prevention, and control analysis.

## Methods

### Ethics statement

The study was approved by the ethical committees of the *Universidade Federal do Amazonas* (CAAE: 97439918.5.1001.5020, Approval number: 2.961.307) following the principles expressed in the Declaration of Helsinki. Written informed consents were obtained from the owners of the street market stores.

### *Trypanosoma cruzi* cultivation

Strains and clones of *Trypanosoma cruzi*, belonging to subpopulations classified between DTUs I to VI: Dm28c (TcI), Y (TcII), INPA 3663 (TcIII), INPA 4167 (TcIV), LL014 (TcV) and CL (TcVI), were obtained from Coleção de Protozoários of Fundação Oswaldo Cruz, Rio de Janeiro, Brazil (Fiocruz, COLPROT, http://www.colprot.fiocruz.br). Epimastigotes were cultured in LIT (Liver Infusion Tryptose - BD, USA) medium supplemented with 10% of heat-inactivated Bovine Fetal Serum (Invitrogen, Massachusetts, USA), at 28 °C for 5 days, to reach logarithmic growth phase. Parasites were harvested by centrifugation (3000x g for 10 minutes, at 4 °C), washed three times with 0.15 M NaCl, 0.01 M phosphate buffer pH 7.2 (PBS) and resuspended in the same solution, prior to artificial contamination of açai pulp samples and DNA extraction. Parasite growth was estimated by counting cells in the Neubauer Chamber hemocytometer using an optical microscope and expressed as parasites/mL.

### Preparation of Guanidine-EDTA Açai (GEA) samples

For analytical validation, açai pulp samples were provided by the Instituto Nacional de Controle de Qualidade em Saúde (INCQS/FIOCRUZ), Rio de Janeiro, Brazil. In addition, 45 samples of açai pulp gathered randomly from different street markets in the city of Coari (Amazonas) in June 2018 were used for the validation of the molecular methodology.

Samples were separated into 5mL aliquots and immediately mixed with an equal volume of a lysis solution containing 6M Guanidine-HCL/0.2 N EDTA pH 8.0 (1:1 ratio). Before DNA extraction, the total volume of açai lysate (GEA) was centrifugated at 10,000 xg for 10 minutes at room temperature. After the centrifugation, GEA supernatants were recovered and stored at 4°C until DNA extraction (Fig 3).

**Fig 3.**
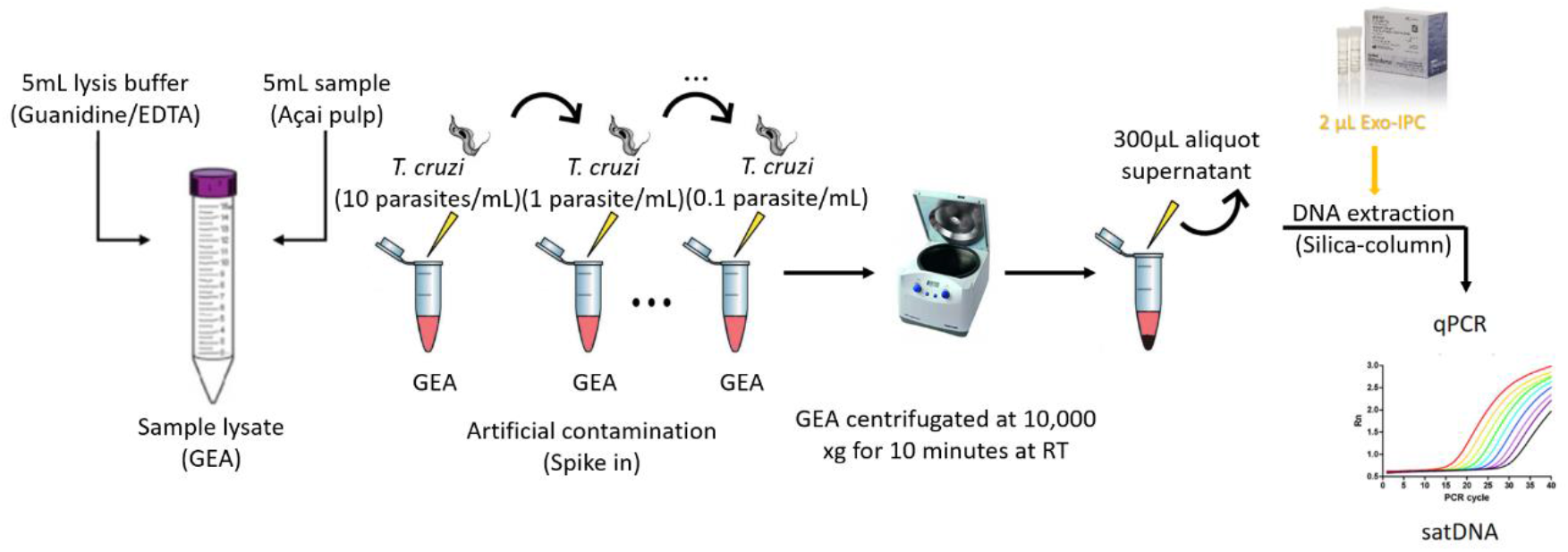
Scheme for the preparation of Guanidine-EDTA Açai (GEA) samples and DNA extraction using the Exogenous Internal Positive Control (Exo-IPC). The scheme shows the DNA extraction from the supernatant of the GEA spiked with EXO-IPC synthetic DNA, using a silica-column based commercial kit.

### DNA extraction

Each 300 μL of GEA supernatant sample was extracted using the High Pure PCR Template Preparation Kit (Roche Life Science, Mannheim, Germany), a kit based on silica-membrane spin columns technology. The DNA purification protocol was carried out according to the manufacturer's instructions with modifications in some steps of the protocol. Briefly, during the proteinase K lysis step, the GEA samples were incubated for 2 hours at 56 °C. In addition, at the DNA elution step, a volume of 100 μL of elution buffer was used to elute the purified DNA. The DNA samples were stored at −20 °C and their purity and concentration were estimated using a NanoDrop 2000c spectrophotometer (Thermo Fisher Scientific, Massachusetts, USA) at 260/280 and 260/320 nm.

In the present study, to monitor the efficiency of the DNA extraction and the absence of inhibitors at PCR, the commercial kit TaqMan Exogenous Internal Positive Control Reagents (Applied Biosystems, Foster City, CA, USA) was used. The EXO-IPC DNA is a synthetic molecule that presents no homology to any DNA sequences available in public databases. Thus, 300 μL GEA supernatant were spiked with 2 μL of the EXO-IPC DNA before DNA extraction. This EXO-IPC is supplied in a commercial kit format that also contains a set of pre-designed primers and TaqMan probe (VIC/TAMRA), targeting the synthetic DNA sequence.

To perform the clinical validation of the methodology, all DNA extractions from the 45 Coari samples were performed in triplicate and DNAs were stored at −20 ◦C until use in qPCR assays.

### Quantitative multiplex real-time PCR (qPCR) assays

Multiplex real time PCR assays were performed for the detection and absolute quantification of the *T. cruzi* DNA in GEA samples. Reactions were carried out in a final volume of 20 μL, containing 5 μL of DNA as a template, 10μL of 2× FastStart TaqMan^®^ Probe Master Mix (Roche Life Science, Mannheim, Germany), 750 nM cruzi1 and cruzi2 primers and 250 nM cruzi3 probe (FAM/NFQ-MGB) targeting *T. cruzi* nuclear satellite DNA (satDNA) and 1μL of the 10× EXO-IPC Mix from the TaqMan Exogenous Internal Positive Control Reagents commercial kit (Applied Biosystems, Foster City, CA, USA), that contains a set of primers and probe (VIC/TAMRA) targeting the synthetic EXO-IPC DNA. Sequences of both sets of primers and probes are described in Table 5. Quantitative assays were performed on the Viia7 equipment (Applied Biosystems, Foster City, CA, USA) with the following cycling conditions: 50 °C for 2 min, 95 °C for 10 min, followed by 45 cycles at 95 °C for 15 s and 58 °C for 1 min. The threshold was set at 0.02 for both targets in all real time PCR assays.

**Table 5.**
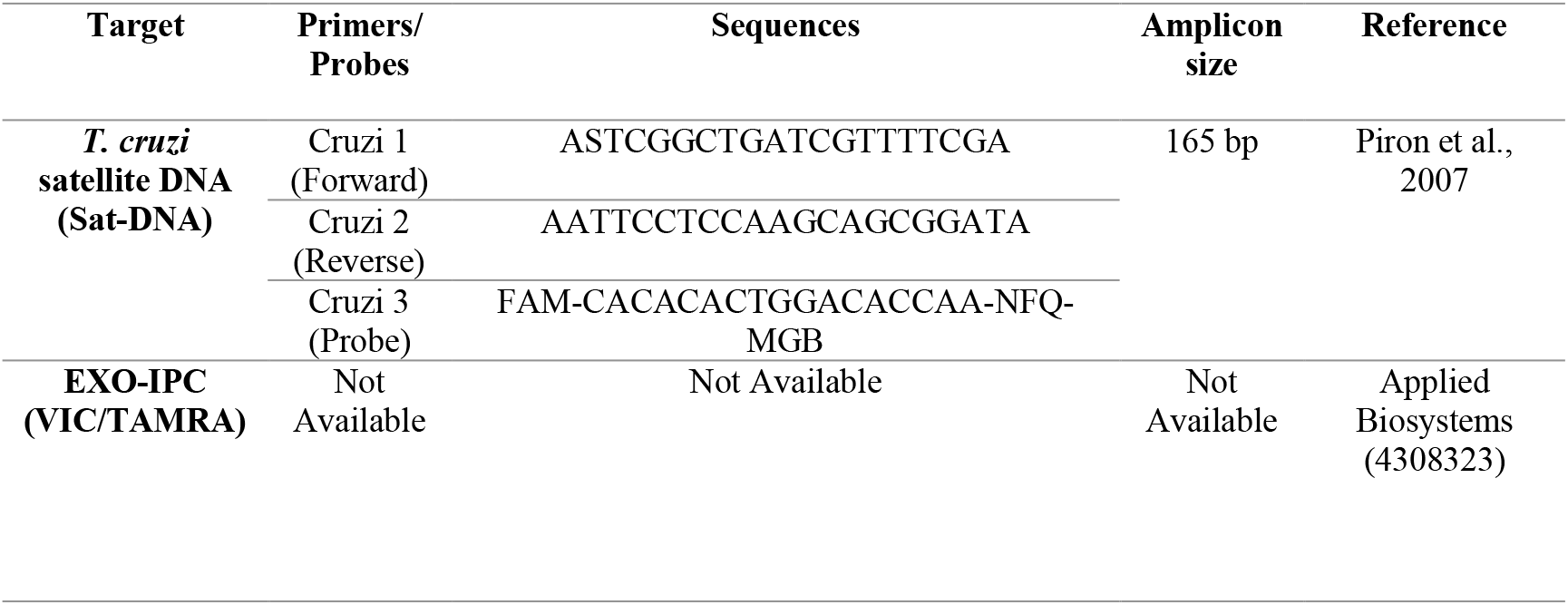
Primer sets and probes sequences for the multiplex qPCR assay.

To build the standard curves for the *T. cruzi* absolute quantification, negative GEA samples were spiked with *T. cruzi* (Dm28c clone, TcI) to reach the 10^6^ epimastigotes/mL concentration, prior to supernatant isolation and DNA extraction. In parallel, DNAs were extracted from negative GEA sample supernatants and pooled, to be used as diluent for the standard curve. The curves were generated by ten-fold serial dilution of DNA from spiked GEA sample in DNA from negative GEA sample, ranging from 10^6^ to 0.1 *T. cruzi* equivalents/mL.

Each 96-well reaction plate included the standard curve, Negative Template Control (ultrapure water instead DNA template) and two positive controls (*T. cruzi* DNA at 10 fg/μL and 1 fg/μL. For each DNA extraction batch (containing up to 11 clinical specimens), one negative control of the DNA extraction step was included, using a negative GEA supernatant sample.

### Inclusivity and exclusivity assays

For the development of qPCR methods some parameters for analytical validation were included, such as: Inclusivity study (i), comprising the detection of the target strains, and exclusivity study (ii), involving the lack of response of closely related non-target strains, which can be potentially cross reactive, but are not expected to be detected [54]. **(i) Inclusivity study:** to assess the ability of this qPCR methodology to detect the *T. cruzi* target, genomic DNA were tested from a representative panel of *T. cruzi* strains/clones belonging to the six different DTUs: Dm28c; Y; INPA 3663; INPA 4167; LL014; CL Brener. Samples were assayed in duplicates, with concentrations ranging from 10^4^ to 10^−1^ parasite equivalents/mL. **(ii) Exclusivity study:** to assess this qPCR methodology’s lack of response from closely related non-target strains, serial dilutions of genomic DNA obtained from other species of trypanosomatids were tested: *Trypanosoma rangeli, Leishmania (Leishmania) amazonensis, Leishmania (Viannia) braziliensis, Herpetomonas muscarum*, and *Chritidia fasciculata*. Samples were assayed in duplicates, with concentrations ranging from 10^4^ to 0.1 parasite equivalents/mL.

### Statistical analyses

All experiments were performed at least in biological triplicates and experimental duplicates and data are reported as arithmetic mean ± standard deviation. All statistical tests were conducted using the Sigmaplot Windows program version 14.0 (Systat Software, Inc., California, USA). Student’s t test or Mann-Whitney Rank-Sum test were adopted to analyze the statistical significance of the apparent differences, according to the parametric or non-parametric distribution of the data. A p-value less than 0.05 was considered statistically significant (p<0.05).

## Acknowledgments

We thank to Instituto Oswaldo Cruz (IOC) and Instituto Nacional de Controle de Qualidade em Saúde (INCQS) for supplying facilities.

